# PanGBank: a large-scale resource of precomputed microbial pangenomes built with PPanGGOLiN

**DOI:** 10.64898/2026.08.05.742796

**Authors:** Jean Mainguy, Téo Lemane, Adelme Bazin, Jérôme Arnoux, Guillaume Gautreau, Claudine Médigue, Alexandra Calteau, David Vallenet

**Author notes:** Jean Mainguy and Téo Lemane contributed equally to this work. Alexandra Calteau and David Vallenet share PI status equally on this work.

## Abstract

PanGBank (https://pangbank.genoscope.cns.fr) is a comprehensive open-access database providing precomputed prokaryotic pangenomes at a broad taxonomic scale. Built upon PPanGGOLiN partitioned pangenome graphs, PanGBank addresses the growing need for large-scale comparative genomics through a standardized, regularly updated, and fully accessible resource. The initial release comprises two complementary collections covering more than 4,600 prokaryotic species from the Genome Taxonomy Database (GTDB), encompassing over 393,000 genomes: GTDB all, maximizing taxonomic and environmental diversity through the inclusion of MAGs and SAGs, and GTDB refseq, focusing on high-quality, annotation-rich genomes. Each species-level pangenome integrates graph-based statistical partitions into persistent, shell, and cloud gene families, together with regions of genomic plasticity (panRGP) and co-localized functional modules (panModule). PanGBank offers multiple access modes, including a REST API, a command-line interface (PanGBank-cli), and an interactive web interface. By combining large-scale pangenome resources with advanced graph-based analyses, PanGBank provides a scalable framework for exploring microbial diversity, genome evolution, functional variation, and the dissemination of adaptive traits across prokaryotic populations, as illustrated by a use case on Acinetobacter baumannii pangenome investigating the distribution and evolution of antimicrobial resistance determinants.

**Graphical Abstract:** 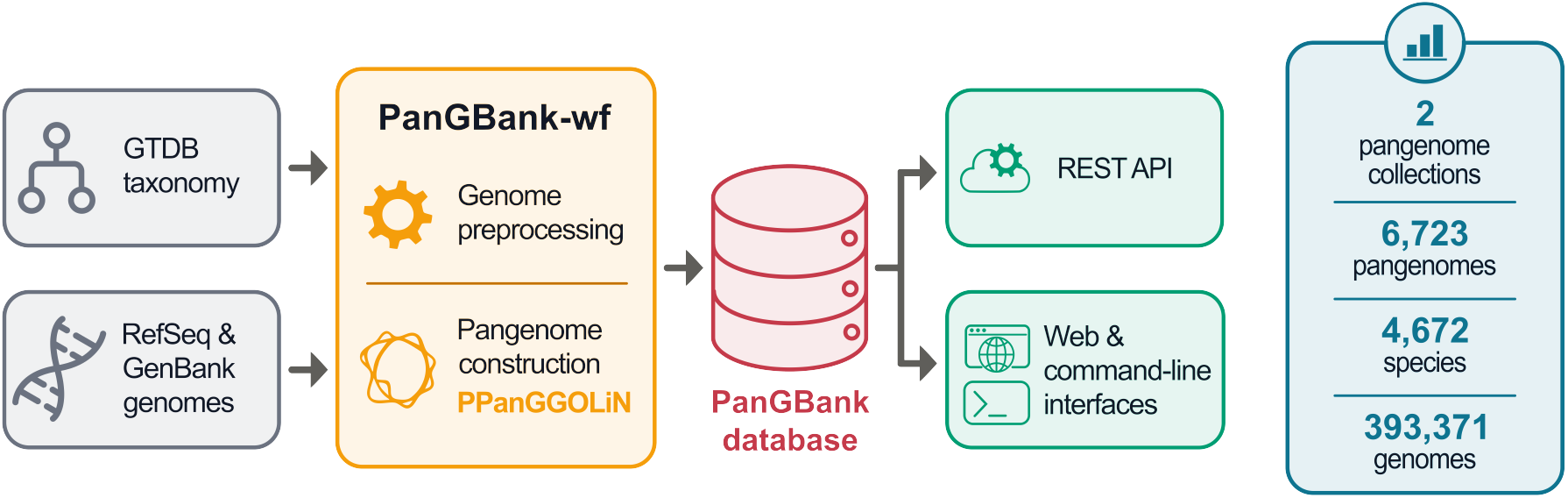

## Introduction

The exponential growth in microbial genome sequencing has fundamentally reshaped the field of prokaryotic biology. NCBI now hosts more than 516,000 prokaryotic genome assemblies in RefSeq, while GenBank contains over 3.3 million (July 2026). To support comparative analyses of this ever-growing collection of sequences, a rich ecosystem of specialized genome databases has been developed, each addressing different analytical needs. Several resources combine large-scale genome collections with integrated annotation and analysis services, such as: the Integrated Microbial Genomes and Microbiomes system (IMG/M, JGI) (1), the Bacterial and Viral Bioinformatics Resource Center (BV-BRC, NIAID) (2), or the MicroScope platform, which further emphasizes synteny-based comparative genomics and expert manual curation (3). Others prioritize exhaustive genome coverage combined with standardized quality control and annotation, like AllTheBacteria, which uniformly assembles all publicly available bacterial isolate sequencing data into currently over 2.7 million genome assemblies (4), or ProGenomes, which provides nearly 2 million high-quality prokaryotic genomes from more than 30,000 species with systematic annotations of mobile genetic elements, biosynthetic gene clusters and habitat metadata (5). Together, these resources have greatly advanced genome-centric comparative microbiology. However, they primarily represent each species through individual genome assemblies, providing only limited support for the analysis of intraspecific diversity.

The concept of the pangenome was motivated by the observation that a single reference genome sequence does not reflect genetic variability within a bacterial species (6). The pangenome was defined as the complete set of genes found across all strains of a species, divided into a core genome

— shared by all isolates — and a dispensable genome — present only in subsets of strains. This framework rapidly became a central paradigm in microbial genomics, as many species were found to harbor open pangenomes that expand continuously with newly sequenced strains, reflecting the pivotal role of horizontal gene transfer (HGT) in prokaryotic evolution. Genomic regions acquired through HGT frequently encode clinically and ecologically relevant functions, including virulence factors, antimicrobial resistance (AMR) determinants, and niche-specific metabolic capabilities, making pangenome analysis of direct relevance to infectious disease research and environmental studies.

The availability of large genomic datasets has driven the development of a growing suite of pangenome inference tools, from early approaches based on “all-against-all” protein comparisons such as Roary (7) and its successor, Panaroo (8), addressing annotation robustness, to pipelines integrating genome downloading, annotation and pangenome construction into unified workflows like PanACoTA (9), or offering graph-based data structures and multi-species scalability such as PanTools (10). Among these, PPanGGOLiN (11; 12) stands out for its statistical graph-based model to classify gene families into *persistent, shell*, and *cloud* genomes rather than relying on arbitrary frequency thresholds. The *persistent* component corresponds to highly conserved gene families shared across most genomes, whereas the *shell* and *cloud* components represent the variable fraction of the pangenome, comprising genes with intermediate and low prevalence, respectively, that contribute to genome plasticity and adaptation. PPanGGOLiN scales to tens of thousands of genomes, including environmental genomes such as metagenome-assembled genomes (MAGs) and single-cell amplified genomes (SAGs), and is further enriched by two complementary modules: panRGP, which predicts Regions of Genomic Plasticity (RGPs) and their insertion hotspots (13), and panModule, which identifies modules of co-occurring and co-localized variable genes within the pangenome graph (14). Together, they are providing a multi-scale view of pangenome structure and genomic plasticity.

Beyond these tools primarily designed for bioinformaticians, several platforms have sought to make pangenome reconstruction and exploration accessible to a broader range of users. PanExplorer (15) provides a web application that performs on-the-fly pangenome analyses using Roary or PanACoTA on user-selected sets of genomes identified by GenBank accession numbers. It offers a suite of interactive visualization modules including: presence/absence matrices, Circos representations, and synteny exploration, together with COG functional annotations, making pangenome analysis accessible to users without bioinformatics expertise. More recently, integrated pipelines such as the one described by Braganza *et al*. (16) have automated the multi-step workflow of genome annotation, gene family clustering, and pangenome generation, using PPanGGOLiN as the core clustering engine and further lowering the barrier to pangenome analysis for users without advanced bioinformatics skills.

Despite these advances in making pangenome reconstruction more accessible, dedicated web resources offering precomputed microbial pangenomes at a broad taxonomic scale remain scarce. panX was among the first, coupling automated pangenome computation with an emphasis on phylogenetic analysis of orthologous gene clusters and interactive visualization; it currently hosts pangenomes for approximately 264 bacterial species (17). ProPan subsequently expanded the taxonomic scope to 1,504 prokaryotic species across 51,882 strains, offering analytical features targeting metabolic and resistance gene dynamics (18). Published more recently, the PanKB resource provides a pangenome knowledgebase encompassing 51 pangenomes from 8 industrially relevant microbial families, comprising 8,402 genomes. It further integrates alleleomic analyses describing intraspecies sequence variation at nucleotide resolution across more than 7 million variants (19). panX and ProPan have not been updated since their initial release and, moreover, neither allows users to download the underlying pangenome data for downstream analyses. Precomputed core and pan gene sets can also be retrieved from orthology-based platforms such as EDGAR (20), whose public section covers 24,317 genomes across 749 mostly genus-level projects, and MBGD (21), whose hierarchical ortholog tables span 1,812 genus-level and 6,268 species-level pangenomes. Both, however, remain restricted to curated sets of complete or near-complete genomes, limiting their applicability to the broader diversity of genomic data now available and underscoring the need for a resource combining broad taxonomic coverage, regular updates, and full data accessibility.

Here we present PanGBank (https://pangbank.genoscope.cns.fr), a freely accessible web database designed to address this gap. PanGBank provides precomputed PPanGGOLiN pangenomes for over 4,600 prokaryotic species derived from two public genome collections sourced from NCBI GenBank and RefSeq, and organized according to Genome Taxonomy Database (GTDB) species clusters to ensure taxonomic consistency (22). For each species, partitioned pangenome graphs together with panRGP predictions and panModule annotations are freely available through an intuitive web interface and a REST API. By tightly integrating PPanGGOLiN workflows with a large set of genomes, PanGBank establishes a unified ecosystem in which precomputed pangenomes can be seamlessly explored, downloaded, and incorporated into downstream analyses. It provides a reference resource for in-depth comparative genomics, enabling users to compare newly sequenced genomes with existing species pangenomes and to contextualize their own genomic datasets within a comprehensive evolutionary framework. By providing partitioned pangenome graphs, genomic plasticity predictions, conserved module annotations, and the complete underlying pangenome data across a broad taxonomic range, PanGBank offers a unique resource for comparative microbial genomics, evolutionary studies, and the analysis of prokaryotic genome dynamics.

### PanGBank overview

PanGBank is an open web resource providing precomputed microbial pangenomes constructed using PPanGGOLiN (11; 12), together with their associated genomic features, at a broad taxonomic scale. Its modular workflow is designed to process any set of genomes associated with a defined taxonomy, allowing multiple pangenome collections to be generated from different genome sources and taxonomic classifications (Fig. 1). As a result, each collection can reflect distinct choices regarding genome sources, inclusion criteria, and species delineation.

**Figure 1.**
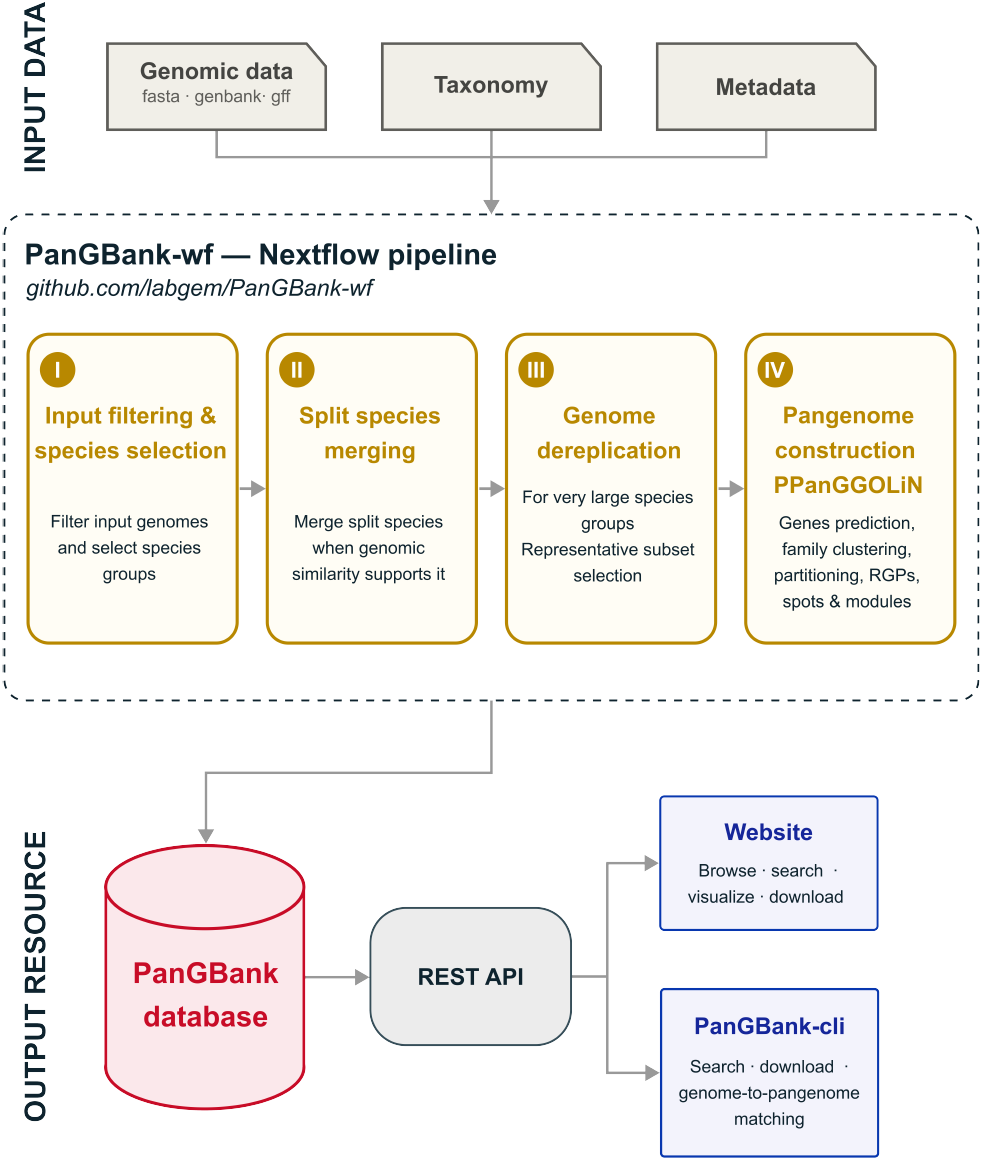
Pangenome construction workflow of PanGBank. Overview of the PanGBank workflow used to generate a collection of pangenomes at the species-level from genomic data and a taxonomy, with optional metadata integration. Input genomes are processed through four main steps within the PanGBank-wf Nextflow pipeline: input filtering and species-group selection, merging of split species when genomic similarity supports it, dereplication of very large species groups to obtain representative genome subsets, and pangenome construction with PPanGGOLiN. The resulting pangenomes are integrated into the PanGBank database, which can be queried through a REST API and accessed via the PanGBank website or a command-line interface.

The initial PanGBank release provides two pangenome collections built from public genomes retrieved from NCBI GenBank and/or RefSeq, together covering more than 4,600 prokaryotic species. Species delineation relies on the GTDB, which provides a phylogenetically consistent and rank-normalized genome-based taxonomy for prokaryotic species (22). By adopting the GTDB species clusters, PanGBank ensures that pangenomes are built from taxonomically coherent genome sets, independent of the classification inconsistencies that can affect public genome repositories. The PanGBank database gathers statistically partitioned pangenome graphs constructed with PPanGGOLiN, along with associated RGP and insertion spot predictions generated by panRGP (13) and conserved module identification from panModule (14). All results are freely accessible through an interactive web interface accessible without registration, as well as through a dedicated REST API that enables programmatic access and seamless integration with PPanGGOLiN command-line workflows (Fig. 1, see ‘PanGBank Access’ section for further details).

### PanGBank workflow

PanGBank is organized into collections of species-level pangenomes. Each collection is constructed from a set of genome sequences with associated taxonomic information using an automated workflow implemented in Nextflow (23) and developed following nf-core guidelines (24). This modular workflow, named PanGBank-wf, is designed to process any set of genomes associated with a defined taxonomy, enabling the generation of pangenome collections from diverse genome sources and classification schemes (Fig. 1). It consists of four main steps: (i) input filtering and species selection, (ii) merging of split species when GTDB is used as the reference taxonomy, (iii) genome dereplication for very large species groups, and (iv) pangenome construction with PPanGGOLiN. Two input files are required: a genome list file specifying genome identifiers and the corresponding file paths to assemblies in FASTA, GenBank or GFF format, and a taxonomy file containing the complete taxonomic assignment for each genome. An additional metadata file can be provided to give additional information about genomes (e.g. strain name, assembly quality metrics, genome completion and contamination). These metadata can be used during the optional filtering step and are propagated to the final pangenome files for downstream analyses (see ‘PanGBank use case’ section for further details). At the end of the workflow, a MultiQC report is generated and available on the PanGBank web site, providing general statistics together with the versions of all software used, thereby ensuring reproducibility.

#### Input filtering and species selection

This step enables optional genome filtering based on quality metrics provided in the metadata input file and then determines the list of species for which pangenomes will be constructed. Only species with at least 15 genomes are retained for pangenome construction, following the minimum number of genomes recommended for PPanGGOLiN. For PanGBank collections based on GTDB, genome completeness values of CheckM v1 (marker-gene based) (25) and CheckM v2 (machine-learning based) (26) are used for filtering. For each genome, the maximum of the two values is used as the completeness score. Filtering is applied at two levels: genomes with completeness *<* 70% are removed, and species are excluded when their GTDB representative genome has completeness *<* 85%. This filtering process aims to prevent the reconstruction of pangenomes from highly incomplete genomes and to reduce potential biases introduced by low-quality environmental genomes, such as MAGs or SAGs.

#### Merging split species in GTDB

This step was specifically designed to handle particular cases when the GTDB taxonomy is used as the reference taxonomic classification. Indeed, GTDB defines species clusters by assigning genomes to a representative genome using average nucleotide identity (ANI) and alignment fraction (AF) thresholds (ANI *≥* 95% and AF *≥* 50%), rather than through all-versus-all comparison of genomes. This representative-centric method is prone to a classical clustering artifact: two genomes can end up in different species clusters despite being more similar to each other than to their own cluster’s representative. To mitigate this issue, we merge closely related species into a single group when they are sufficiently close at the genomic level.

Identifying all such cases would require pairwise ANI/AF comparisons across all GTDB species, which is not tractable at this scale. We therefore restrict this correction to species sharing the same base name and differing only by an alphabetic suffix (e.g. s Proteus vulgaris, s Proteus vulgaris B and s Proteus vulgaris C), a naming convention used in GTDB to resolve polyphyletic groups in its reference tree, which often corresponds to species that remain close in terms of ANI and AF. For each group of GTDB species sharing the same base name, mean ANI and AF values are computed between all species pairs using Skani (27). A graph is then constructed in which an edge connects two species when their mean ANI exceeds 95% and their mean AF exceeds 50%, following GTDB species delineation criteria. Connected components in this graph define the merged species groups used in downstream pangenome construction steps.

#### Genome dereplication

To facilitate the distribution and subsequent analysis of pangenomes while limiting storage requirements and computational resources, species-level pangenomes are restricted to a maximum of 2,000 genomes. For species exceeding this threshold, genomes are dereplicated using a tree-based clustering strategy. Pairwise genomic distances are first computed with Mash (28), and a Neighbor-Joining tree is inferred from the resulting distance matrix using Quicktree (29). The tree is then partitioned into 2,000 clusters, from which a single representative genome is selected based on the highest completeness estimate (the maximum value of the CheckM v1 and CheckM v2 completeness scores), with assembly quality metrics used to resolve ties when necessary. Finally, any NCBI reference genomes and GTDB representative genomes missing from the selection are reintroduced to ensure that important reference data are retained.

#### Pangenome creation

For each species or species-group cluster, the complete PPanGGOLiN workflow (*all* subcommand) is executed using all the corresponding genomes as input to generate a partitioned pangenome graph, in which gene families are classified into *persistent, shell*, and *cloud* components. The resulting pangenome includes the prediction of RGPs, their insertion spots, and conserved modules composed of co-occurring and co-localized genes within the variable genome. Genome metadata provided as input to the workflow are also integrated into the final pangenome files for downstream analyses. In addition, a Mash sketch is built from *persistent*-family sequences of each pangenome, enabling rapid identification of the pangenome most closely related to a query genome using the *match-pangenome* command of the PanGBank command-line interface (CLI) tool (see ‘PanGBank Access’ and ‘PanGBank use case’ sections for further details).

### PanGBank content

The first collections made available in the PanGBank resource were constructed from the GTDB release 232, comprising GenBank and RefSeq genomes selected according to GTDB quality criteria. Two complementary collections were generated: GTDB all and GTDB refseq. Genomes were retrieved from NCBI using the ncbi-datasets-cli tool (30), based on their assembly accession. The GTDB all collection includes genomes from both GenBank and RefSeq, downloaded in FASTA format, whereas GTDB refseq is restricted to RefSeq genomes, which were downloaded in GenBank format in order to preserve the original gene predictions and functional annotations. This design allows GTDB all to maximize genome sampling and taxonomic coverage, while GTDB refseq provides high-quality, annotation-rich pangenomes that are better suited to functional analyses.

The GTDB all collection covers more than 4,600 prokaryotic species, compared with 2,044 species in GTDB refseq. This broader taxonomic coverage results in substantially higher numbers of RGPs, modules and gene families, reflecting the greater genomic diversity represented in GTDB all (Table 1). Despite these differences in scale, both collections are dominated by relatively small pangenomes comprising fewer than 50 genomes, while only about one hundred species are represented by more than 1,000 genomes (Fig. 2). Among these highly sampled taxa, a limited number required dereplication to reduce redundancy—24 species in GTDB refseq and 27 in GTDB all.

**Table 1.** Description of the content of the PanGBank collections based on the GTDB taxonomy release 232.

| Collection name and version | Pangenomes | Genomes | Pangenomes with merged species | Pangenomes with genome dereplication | RGPs | Modules | Families |
| --- | --- | --- | --- | --- | --- | --- | --- |
| GTDB_all 2.1.0 | 4,679 | 382,760 | 148 | 27 | 14,389,864 | 1,010,214 | 40,239,843 |
| GTDB_refseq 2.0.0 | 2,044 | 219,611 | 127 | 24 | 8,706,710 | 579,670 | 19,803,205 |

**Figure 2.**
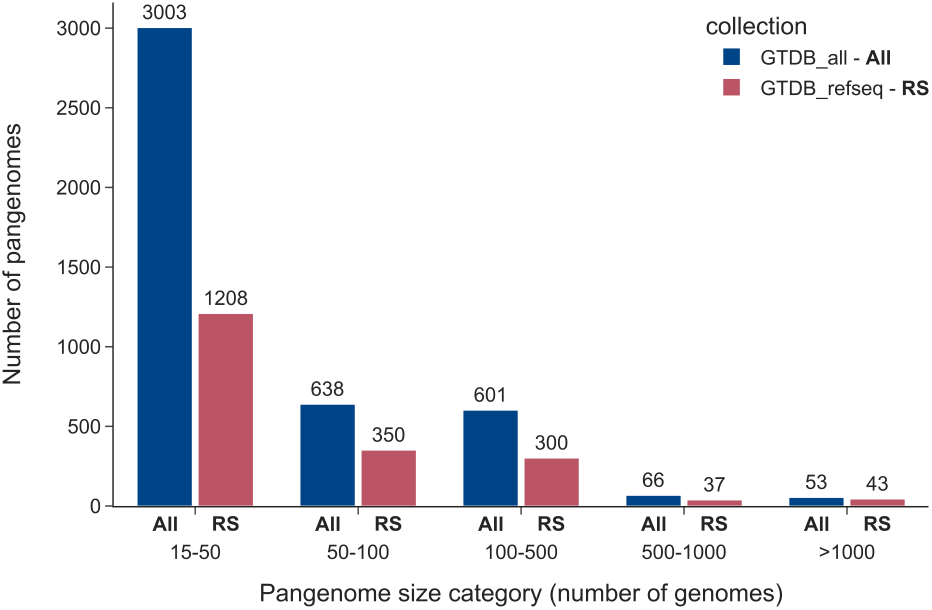
Distribution of the number of genomes per pangenome in the GTDB all (All) and GTDB refseq (RS) collections. Pangenomes were grouped according to the number of genomes included in each pangenome. Numbers above bars indicate the number of pangenomes in each category.

These species mainly correspond to bacterial taxa of major public health importance, which are overrepresented in public genome databases.

As shown in Figure 3A, approximately 40% of the genomes included in GTDB all are MAGs or SAGs. By contrast, GTDB refseq consists almost exclusively of genomes from isolates. Both collections are dominated by bacterial pangenomes, and GTDB all contains only about 150 archaeal pangenomes, largely owing to the inclusion of genomes from environmental samples (Fig. 3B). GTDB all has a significantly broader taxonomic coverage, encompassing 55 phyla, of which around 40% are represented exclusively by environmental genomes (Fig. 3C).

**Figure 3.**
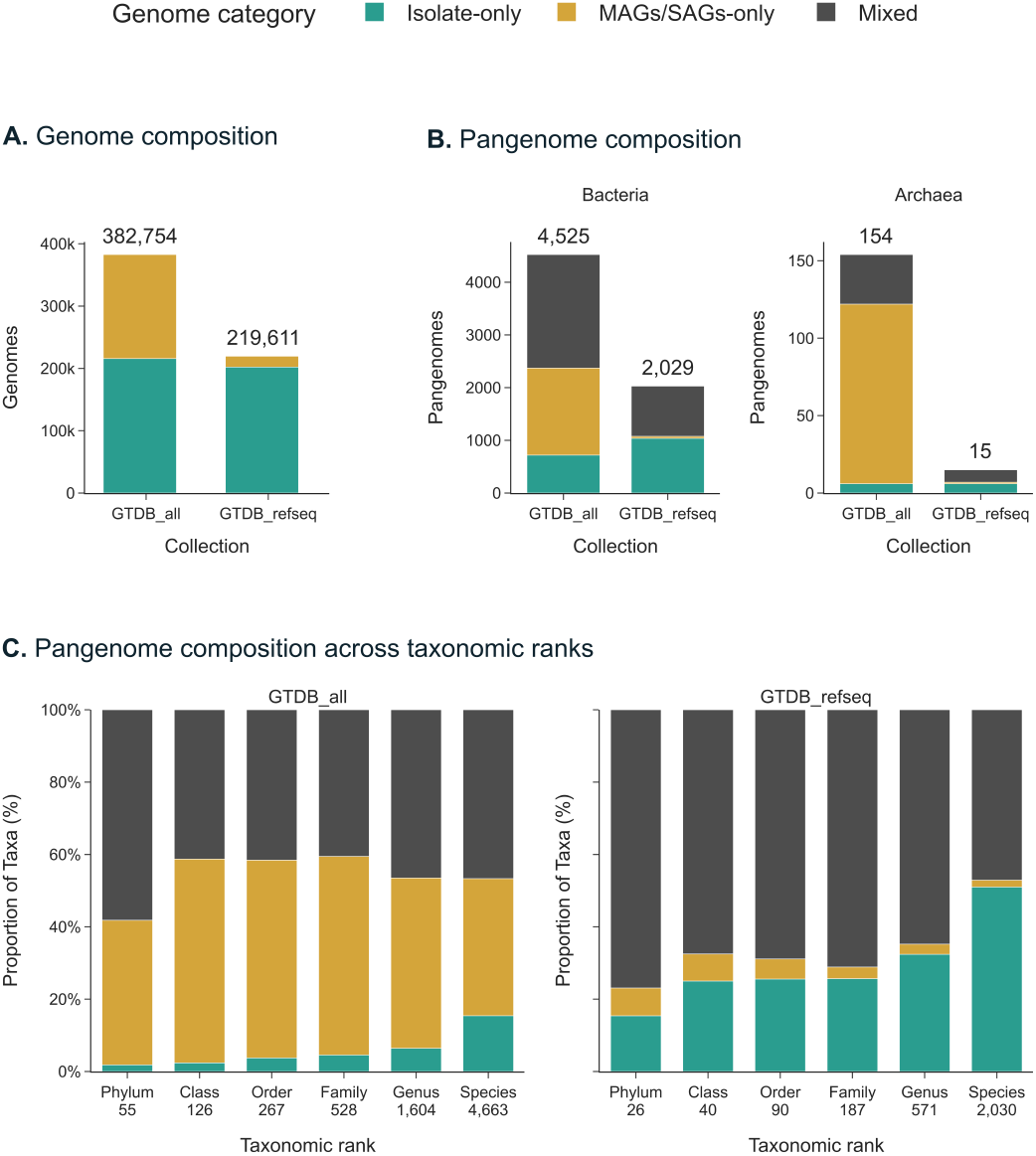
Comparison of PanGBank collections by genome category, and taxonomic rank. Pangenomes were classified according to the type of genomes they contain: Isolate-only (containing only genomes from isolates), MAG/SAG-only (containing only MAGs and/or SAGs), or Mixed (containing both genomes from isolates and MAGs/SAGs). (A) Total number of genomes in each collection (GTDB all and GTDB refseq) grouped by genome category. (B) Number of pangenomes in each collection by domain (Bacteria and Archaea) and genome category. (C) Proportion of pangenomes by genome category across taxonomic ranks (from phylum to species).

The genome-quality selection criteria applied during the reconstruction of both collections ensured that only high-quality MAGs and SAGs were retained, reducing potential biases associated with incomplete or fragmented genomes during pangenome partitioning. This is reflected by the strong agreement in the proportions of *persistent, shell*, and *cloud* genomes observed in GTDB all and GTDB refseq (Supplementary Fig. S1 and Fig. S2), despite a slight shift toward lower *persistent* fractions in some GTDB all pangenomes (the median decrease across matched species was only 0.4 percentage points).

### PanGBank access

PanGBank provides three complementary interfaces for accessing its pangenome resources: a REST API, an interactive web interface and a command-line interface (CLI).

The REST API (https://pangbank-api.genoscope.cns.fr) serves as the primary access layer, exposing collections, pangenomes, genomes, and associated metadata through programmatic endpoints. It provides the backend for both the web interface and the CLI, enabling integration of PanGBank resources into computational workflows. The API also ships a companion Python SDK that exposes the same functionality as the REST API through a pure Python interface.

The web interface (https://pangbank.genoscope.cns.fr) enables users to browse, search, and visualize pangenome data across collections and taxonomy (Fig. 4). This interface was primarily designed for use on desktop and laptop computers, providing an optimal user experience on large screens, while remaining functional and responsive on tablets and smartphones.

**Figure 4.**
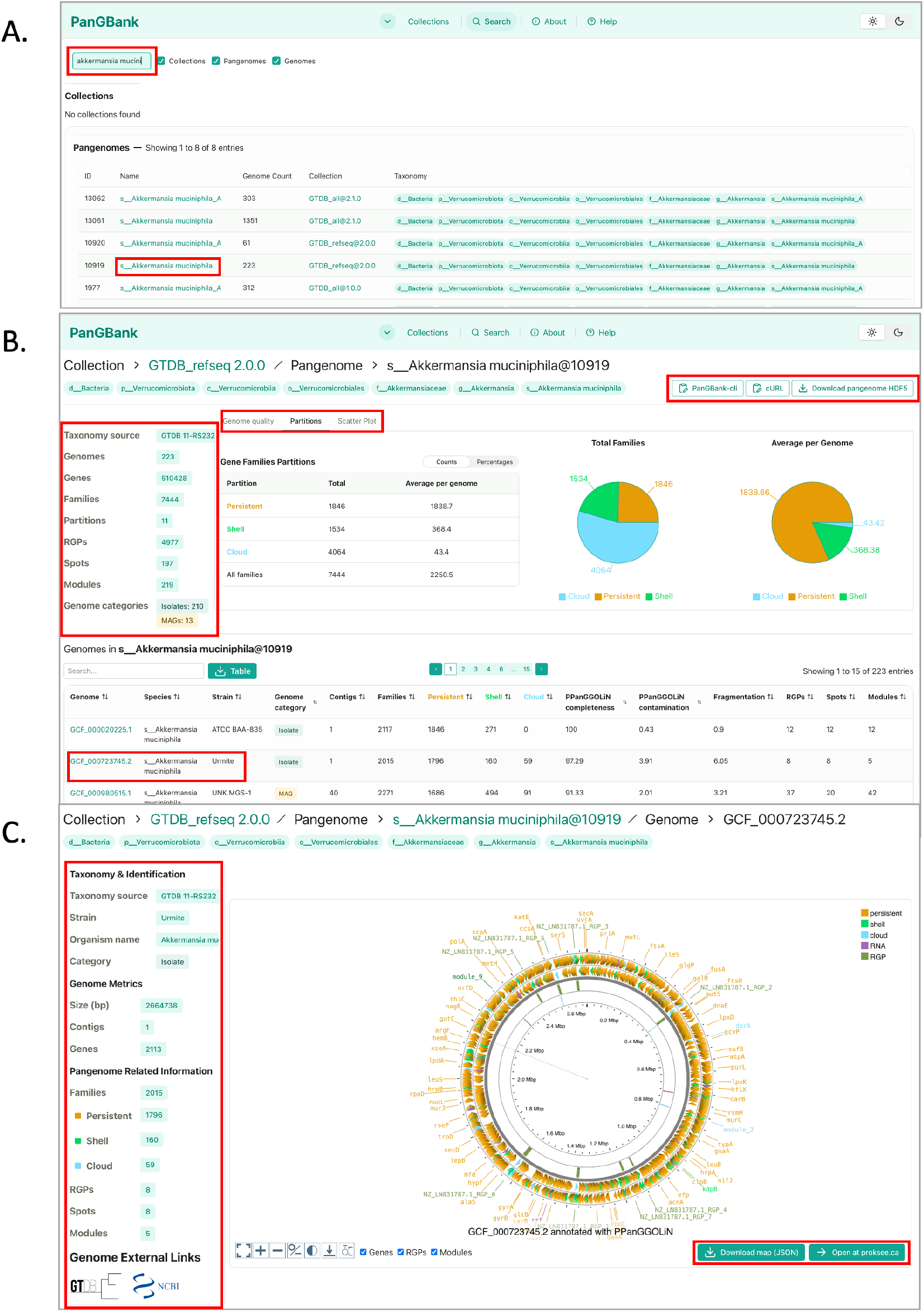
Overview of PanGBank web interface. (A) The search page enables users to query keywords across collections, pangenomes, or genomes, and provides direct links to the corresponding dedicated pages. (B) The pangenome page allows users to access the full set of metrics computed for a given pangenome, as well as the complete list of genomes it comprises. This page features interactive and customizable visualizations, including genome quality plots, partition distributions, and scatter plots, allowing researchers to explore the data according to their specific needs. At the bottom of the page, a detailed table lists all genomes composing the pangenome together with their associated metrics. From this page, users can directly download the pangenome as an HDF5 file or retrieve ready-to-use commands to obtain it via the PanGBank CLI or cURL. (C) The genome page presents pangenome-derived metrics at the individual genome level and provides direct links to the corresponding GTDB and NCBI reference pages. It offers genome-level visualization through interactive CGView maps enriched with genome and pangenome annotations—including partitions, regions of genomic plasticity (RGPs), and modules—and, where available, provides access to detailed gene annotations. Genome maps can be downloaded as JSON files or be opened with ProkSee, allowing users to further explore and customize the visualizations outside the platform.

It supports multi-scale exploration with interactive tables and visualizations, ranging from collection-level summaries comparing pangenome metrics across species, to detailed pangenome pages providing statistics on gene families, RGPs, spots and modules for each genome. At the genome level, interactive visualization is provided through a CGView map (31) enriched with genome and pangenome annotations.

The PanGBank CLI (https://github.com/labgem/PanGBank-cli; distributed through PyPI and Bioconda) provides programmatic access to the same resources for automated analyses. Users can search pangenomes by taxon or genome identifier, retrieve metadata and summary statistics, and download PPanGGOLiN pangenome files in HDF5 (Hierarchical Data Format version 5) for downstream analyses. The CLI additionally supports genome-to-pangenome matching using Mash sketches computed from *persistent* gene families, enabling rapid identification of the most suitable reference pangenome for an input genome. Once retrieved, the corresponding pangenome can be directly used with PPanGGOLiN to project pangenome annotations onto the query genome.

### PanGBank use case

To illustrate the potential of PanGBank for large-scale comparative genomics, we analyzed the pangenome of *Acinetobacter baumannii*, a major opportunistic pathogen characterized by extensive genome plasticity and the rapid acquisition of antibiotic resistance determinants. To ensure reproducibility and facilitate the adaptation of this analysis to other datasets, all analyses presented in this use case are provided as a Jupyter notebook (see ‘Data availability’ section). The PanGBank CLI was used to retrieve pangenome data, while the latest PPanGGOLiN functionalities (12) were used to integrate metadata, project pangenome annotations onto an external genome sequence and cluster RGPs according to their gene family content.

First, the distribution of AMR genes was investigated across the entire *A. baumannii* pangenome. The corresponding pangenome was retrieved from the GTDB refseq collection v2.0.0 containing a total of 2,002 genomes. Representative sequences from all pangenome gene families were exhaustively screened for AMR determinants using AMRFinderPlus v4.2.7 (database version 2026-05-15.1) (32). The resulting AMR annotations were then integrated as metadata into the pangenome files, enabling their exploration alongside pangenome structure and gene family classifications. A total of 123 pangenome gene families were associated with AMR (i.e. 79 assigned to the *cloud* genome, 39 to the *shell* genome and only 5 to the *persistent* genome; Supplementary Fig. S3), with half of them involved in beta-lactam or aminoglycoside resistance (Supplementary Fig. S4). The small number of AMR-associated families in the *persistent* genome is consistent with the limited set of intrinsic resistance determinants conserved across *A. baumannii* strains (33), whereas the large accessory fraction highlights the major contribution of variable genomic content to the resistome diversification. In particular, the predominance of AMR-associated gene families in the *cloud* genome suggests that a substantial fraction of the resistome is strain-specific. However, their notable presence in the *shell* genome reveals the existence of a shared accessory resistome circulating within *A. baumannii* populations, likely shaped by horizontal gene transfer events. To further investigate the dynamics of AMR gene dissemination, we extracted all predicted RGPs and their associated insertion spots from the *A. baumannii* pangenome to identify resistance islands and hotspots of insertion (Supplementary Fig. S5). We identified 3 hotspots (spots 7, 11 and 47) present in more than 500 genomes, for which more than 40% of the associated RGPs contained AMR genes. Spot 47 harbored the most diverse repertoire of AMR genes, comprising 33 gene families, including those found within AbaR1 of strain AYE (Supplementary Fig. S6), one of the largest resistance islands identified in *A. baumannii* (34). This resistance hotspot corresponds to the well-known insertion in the chromosomal *comM* gene and exhibits a wide diversity of resistance islands, differing in gene content, size, and AMR gene composition (35). Using the RGP clustering functionality of PPanGGOLiN, we identified 56 clusters of RGPs within this hotspot, sharing 80% of their gene family content (i.e. *max grr* parameter set to 0.8, defined as the number of shared gene families divided by the larger gene family count of the two compared RGPs) (Supplementary Fig. S7). This closely matches the 53 distinct genetic configurations previously described by Bi *et al*., supporting the ability of our approach to capture the diversity of resistance islands within an insertion spot.

Second, we focused our analysis on strain ABO21-A003, which was isolated from a dog during the surveillance of a veterinary intensive care unit (36). The genome of this strain was not included in the *A. baumannii* pangenome of PanGBank GTDB refseq collection. Therefore, the projection functionality of PPanGGOLiN was used to transfer the pangenome annotation onto the genome. Among the predicted RGPs, four contain AMR genes (Supplementary Table S1). Three mapped to previously identified hotspots (spots 7, 11 and 47), whereas the RGP harboring the largest repertoire of AMR genes (CP136181.1 RGP 5) mapped to spot 31 (Supplementary Fig. S8). As already reported by André *et al*., this region corresponds to the AbGRI2 resistance island and contains three copies of *bla*_*TEM1*_ and two of *aphA1* in strain ABO21-A003, involved in beta-lactam and kanamycin resistance. These results demonstrate that our projection approach accurately identifies both the genomic insertion hotspot and the associated complex resistance island architecture in a genome absent from the reference pangenome.

Together, these examples illustrate how PanGBank enables both large-scale exploration of pangenome AMR gene distribution and rapid characterization of resistance islands in newly sequenced genomes through pangenome projection. By combining precomputed pangenomes with PPanGGOLiN functionalities, PanGBank provides an efficient framework for investigating the diversity, evolution and dynamics of the variable genome, including the dissemination of antimicrobial resistance determinants.

## Discussion and future direction

As a whole, PanGBank collections provide microbiologists with a valuable and scalable framework for investigating genomic diversity both within and across species, thereby bridging the gap between large-scale comparative genomics and detailed functional analysis. They can be explored through two complementary tools devoted to the analysis of pangenome graphs: (i) PPanGGOLiN (11; 12) enables pangenome information to be projected onto newly sequenced genomes, thus allowing researchers to rapidly place novel isolates within an existing pangenomic and functional context; (ii) PANORAMA (37) supports comparative analyses of pangenome graphs across multiple species, enabling the identification of conserved and lineage-specific genomic features at a broader evolutionary scale. Together, these tools transform static reference collections into a dynamic and extensible ecosystem for exploring microbial diversity.

The PanGBank GTDB-based collections will be updated with each new GTDB release, ensuring continuous consistency with the latest genomic and taxonomic advances. This regular update cycle is essential given the rapid pace at which new genomes are sequenced and taxonomic assignments are refined; it thus ensures that users are always working with pangenome resources that accurately reflect the current state of microbial systematics and sequencing. Beyond these regular updates, additional collections tailored to specific research projects (e.g. antimicrobial resistance, biotechnological applications) or environmental niches (e.g. host-associated microbiomes, extreme environments) may also be released. By incorporating dedicated metadata and customized genome sets, these collections will extend the applicability of PanGBank to more targeted and applied research questions.

Future developments of the PanGBank workflow will incorporate functional annotation of gene families, enabling the identification of protein families, protein domains, and AMR genes, together with the prediction of biological processes such as metabolic pathways and defense systems through PANORAMA. This enhancement is a critical step forward, as it will enable the collections to move beyond purely structural and taxonomic descriptions of pangenomes towards a more integrated representation, linking the diversity of gene families to their functional context. By providing consistent and systematically annotated functional information across all collections, this development will not only enable users to describe how gene content varies within a single species and between species, but also to interpret the biological and ecological significance of that variation, for instance, by linking specific gene families to metabolic capabilities, virulence, or resistance phenotypes.

In the short term, the pangenomes made available through PanGBank will be further exploited within the MicroScope microbial genome annotation platform (3), providing a direct link between large-scale, standardized pangenome resources and one of the reference environments for comparative and functional genome analysis. This integration will allow MicroScope users to benefit from the pangenomic context when annotating individual genomes, while expanding the range of species that can be analyzed within the platform.

Beyond genome-centric applications, PanGBank pangenomes will also serve as a reference backbone for MetaPanG, a new software tool currently under development for the characterization of metagenomic samples using precomputed pangenomes (Lemane *et al*., in preparation). This tool aims to identify and quantify strains present in a metagenomic sample, along with their associated gene families, while highlighting genes that may remain undetected due to limited sequencing depth. By directly comparing metagenomic reads against PanGBank pangenome graphs, which capture the full genomic diversity of each species, MetaPanG will enable more precise taxonomic and functional profiling of complex microbial communities. These developments will further extend the scope of applications supported by PanGBank, from single-genome and comparative genomics analyses to community-level metagenomic analyses.

## Data availability

PanGBank web user interface is available at https://pangbank.genoscope.cns.fr and is served through a REST API accessible at https://pangbank-api.genoscope.cns.fr. Pangenome data and associated resources are made freely available under the Creative Commons Attribution-ShareAlike 4.0 International (CC BY-SA 4.0) License. The GTDB-based pangenome collections use and adapt taxonomy from the Genome Taxonomy Database (GTDB), which is also distributed under CC BY-SA 4.0.

The PanGBank workflow, API and CLI are released under the CeCILL v2.1 license. Their source code is available on GitHub at https://github.com/labgem/PanGBank-wf, https://github.com/labgem/PanGBank-api and https://github.com/labgem/PanGBank-cli. The versions used in this manuscript are archived in Software Heritage under the following SWHIDs:

~~~
swh:1:dir:601cfe76b616d459447f24cdf276af5c71720322
~~~

for PanGBank-wf v0.1.4,

~~~
swh:1:dir:4323e2bd37c19f642e02badf6718eac9517dd843
~~~

for PanGBank-api v0.5.0,

~~~
swh:1:dir:bf262a5a778e3c2089ab4645aa6c5449b15f2af8
~~~

for PanGBank-cli v0.2.0.

A GitHub repository containing a Jupyter notebook with all the code required to reproduce the use case presented in this manuscript is available at https://github.com/labgem/PanGBank-tutorial. The notebook can be executed locally using Docker or Conda, or directly in Google Colaboratory.

## Author contributions statement

Conceptualization: D.V., A.C., G.G. and A.B. Funding Acquisition: A.C., C.M., G.G., D.V. Supervision: A.C. and D.V. Methodology: J.M., T.L., D.V. Data Curation: A.B., J.A., J.M., T.L. Software: J.M. and T.L. Investigation, Validation, Visualization: J.M., T.L., A.C. and D.V. Writing – original draft: A.C., J.M. and D.V. Writing – review & editing: A.C., D.V., J.M., T.L., C.M., J.A., A.B. and G.G.

## Acknowledgments

The authors thank previous developers who have worked on an early version of this resource: Mathieu Dubois, Rémi Planel, Laura Burlot, Mathieu Gachet and Paul Amours. We also thank Alexandre Protat for his help in designing the database model. Artificial intelligence tools (chatGPT-5.6 Sol and Claude Sonnet 5) were used to improve the clarity and readability of the manuscript, to generate initial draft versions of the figures (Fig. 1 and graphical abstract) and to assist with software development. All outputs were critically reviewed, validated, and edited by the authors, who take full responsibility for the scientific content, analyses, conclusions, and software implementation presented in this work.

