## Supplementary for "PanGBank: a large-scale resource of precomputed microbial pangenomes built with PPanGGOLiN"

**This PDF file includes:**

- Supplementary Table S1
- Supplementary Figures S1, S2, S3, S4, S5, S6, S7 and S8

### Supplementary Table and Figures

**Table S1: AMR gene content of resistance-carrying RGPs in *Acinetobacter baumannii* canine isolate ABO21-A003.** Each row corresponds to one Region of Genomic Plasticity (RGP) identified in the projected genome that carries at least one AMR gene family. Spot IDs refer to insertion spots defined in the *A. baumannii* pangenome. Gene copy numbers are indicated where a family is present more than once within the same RGP. CP136181.1_RGP_5, mapping to spot_31, harbours the largest AMR gene load with five genes conferring aminoglycoside and beta-lactam resistance, consistent with the AbGRI2 resistance island structure in strain ABO21-A003.

| RGP | Spot ID | # AMR genes | AMR classes | Genes |
|---|---|---|---|---|
| CP136181.1_RGP_5 | spot_31 | 5 | Aminoglycoside<br>Beta-lactam | $2\times$ *aph(3')-Ia*<br>$3\times$ *bla*$_{TEM\text{-}1}$ |
| CP136181.1_RGP_12 | spot_47 | 3 | Aminoglycoside<br>Tetracycline | *aph(3")-Ib*<br>*aph(6)-Id*<br>*tet(B)* |
| CP136181.1_RGP_0 | spot_11 | 1 | Efflux | *adeC* |
| CP136181.1_RGP_1 | spot_7 | 1 | Fosfomycin | *abaF* |

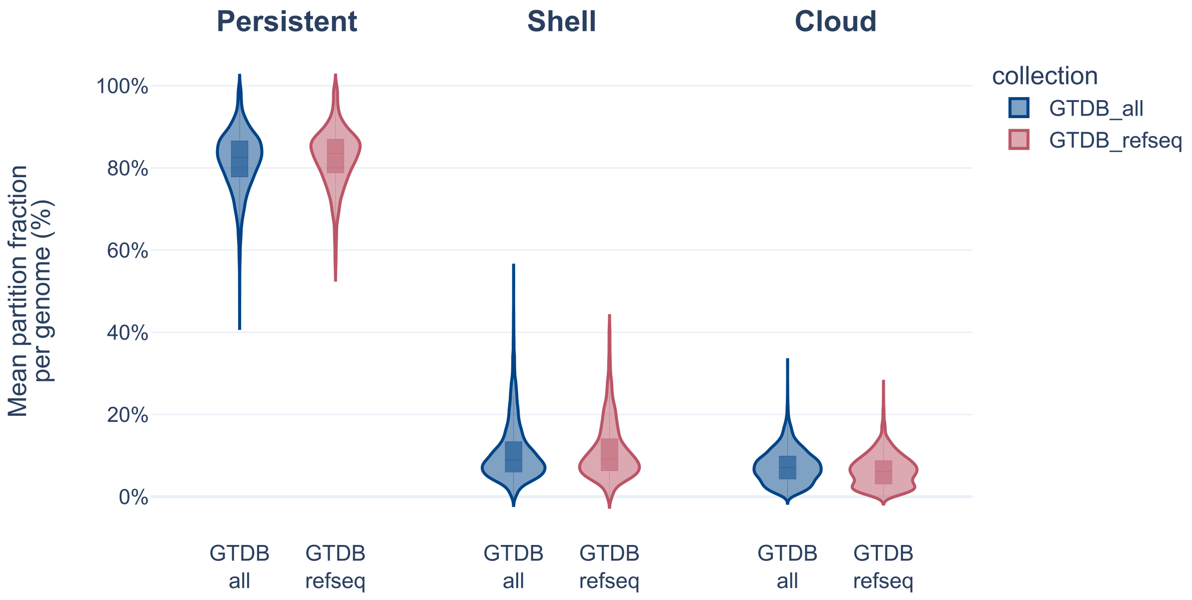


**Figure S1: Distribution of the mean partition fraction per genome across pangenomes.** For each pangenome, the fractions of gene families assigned to the *persistent*, *shell*, and *cloud* partitions were calculated for each genome and then averaged across all genomes in the pangenome. Violin plots show the distributions of these partition fraction means for the GTDB_refseq and GTDB_all collections.

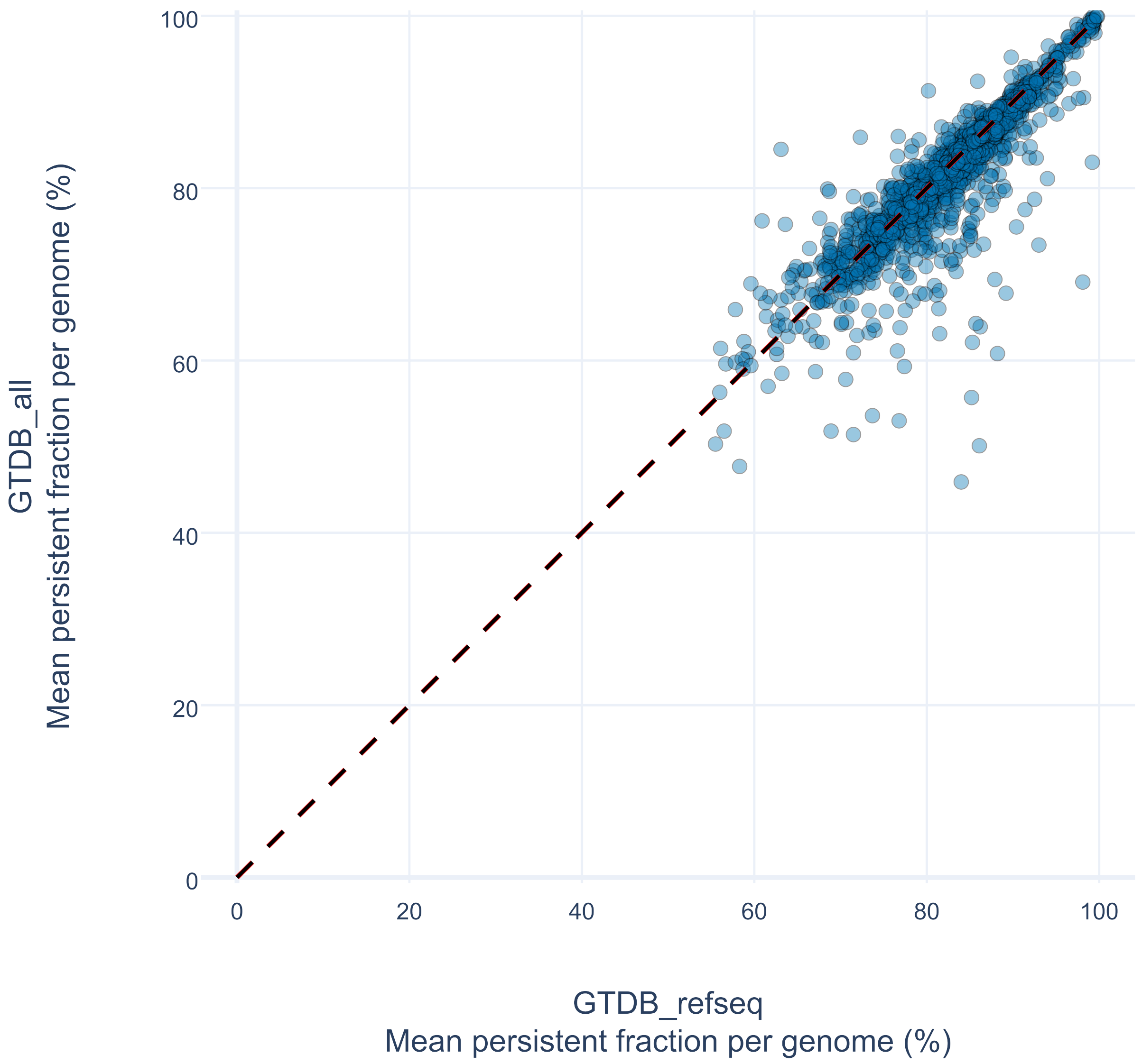


**Figure S2: Comparison of the mean persistent partition fraction per genome between the GTDB_refseq and GTDB_all collections.** Each point represents a species for which a pangenome was computed in both collections. For each pangenome, the fraction of gene families assigned to the persistent partition was calculated for every genome and averaged across all genomes. The dashed line indicates equality between the two collections.

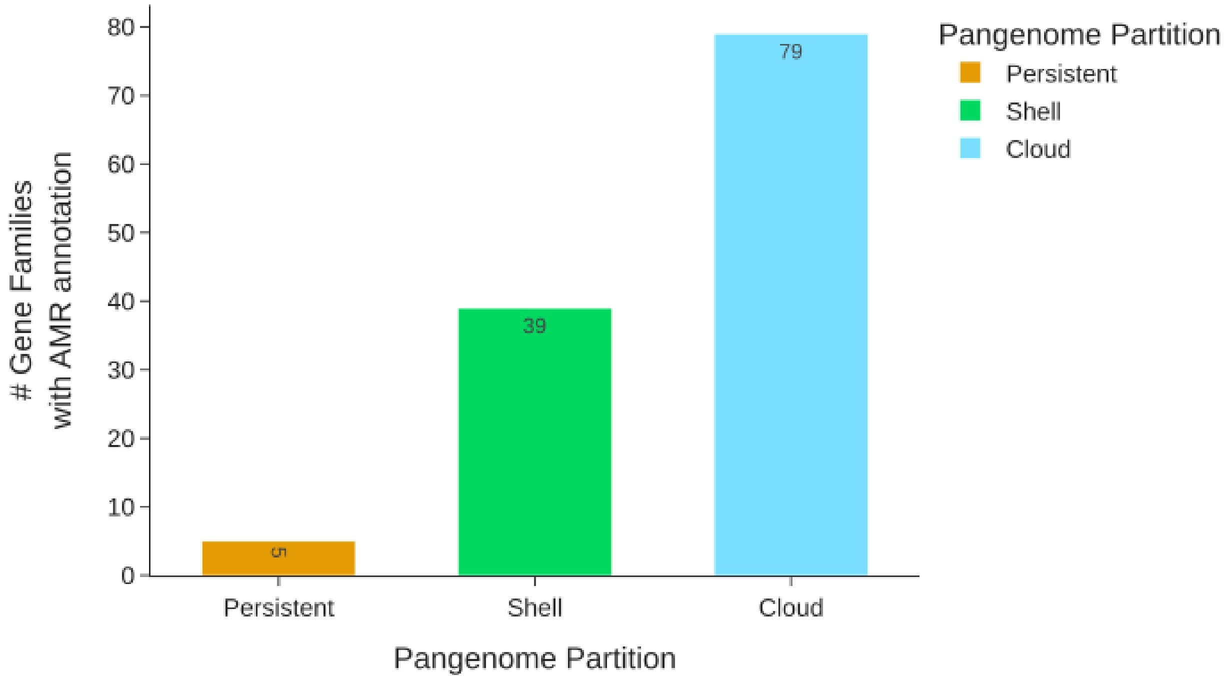


**Figure S3: Distribution of AMR gene families across partitions for the *Acinetobacter baumannii* pangenome.** The majority of AMR gene families are *cloud* (79) or *shell* (39), with only 5 *persistent* families.

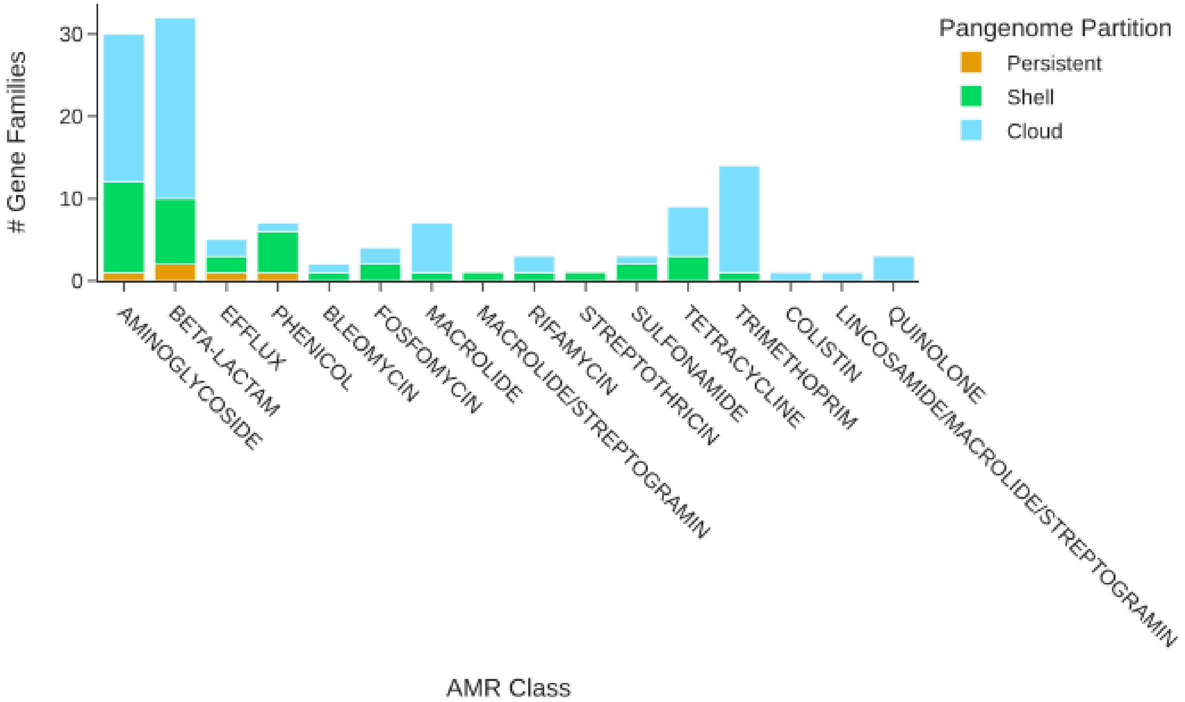


**Figure S4: Distribution of AMR gene families by class and partitions for the *Acinetobacter baumannii* pangenome.** Aminoglycoside and beta-lactam classes contain the highest number of AMR families, most of which belong to the *shell* and *cloud* partitions.

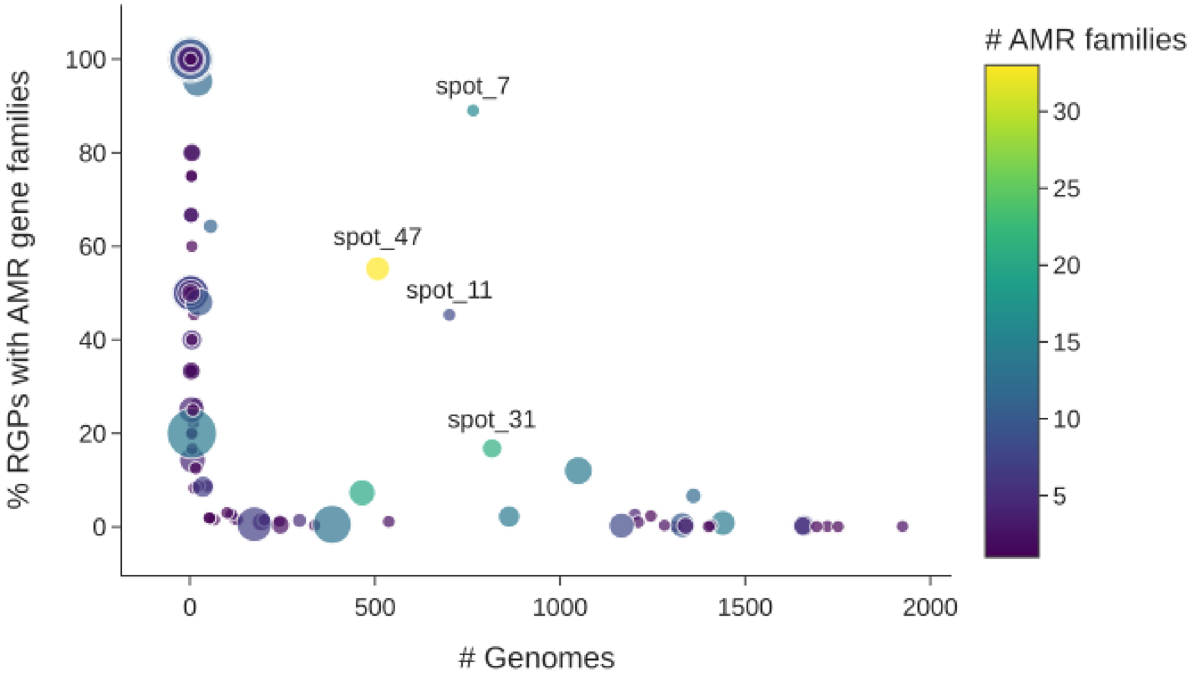


**Figure S5: Distribution and diversity of AMR gene families across insertion spots in the *Acinetobacter baumannii* pangenome.** Each point represents an insertion spot in the pangenome containing at least one AMR gene family. The x-axis indicates the number of genomes carrying an RGP inserted at the corresponding spot; the y-axis shows the percentage of RGPs within each spot that contain at least one AMR gene family. Point size is proportional to the mean number of AMR gene families per RGP, and the color scale indicates the number of distinct AMR gene families identified at each spot (from low in purple to high in yellow). Spots 7, 11, and 47 represent major AMR hotspots, combining high prevalence across genomes ($> 500$) with a high proportion of AMR-carrying RGPs ($> 40\%$). Spot_47 exhibits the largest AMR repertoire, with 33 AMR gene families. Spot_31, although containing fewer AMR-associated RGPs, harbors a highly diverse AMR repertoire comprising 21 gene families. Spot_47 and Spot_31 correspond to the AbaR and AbGRI2 resistance islands, respectively.

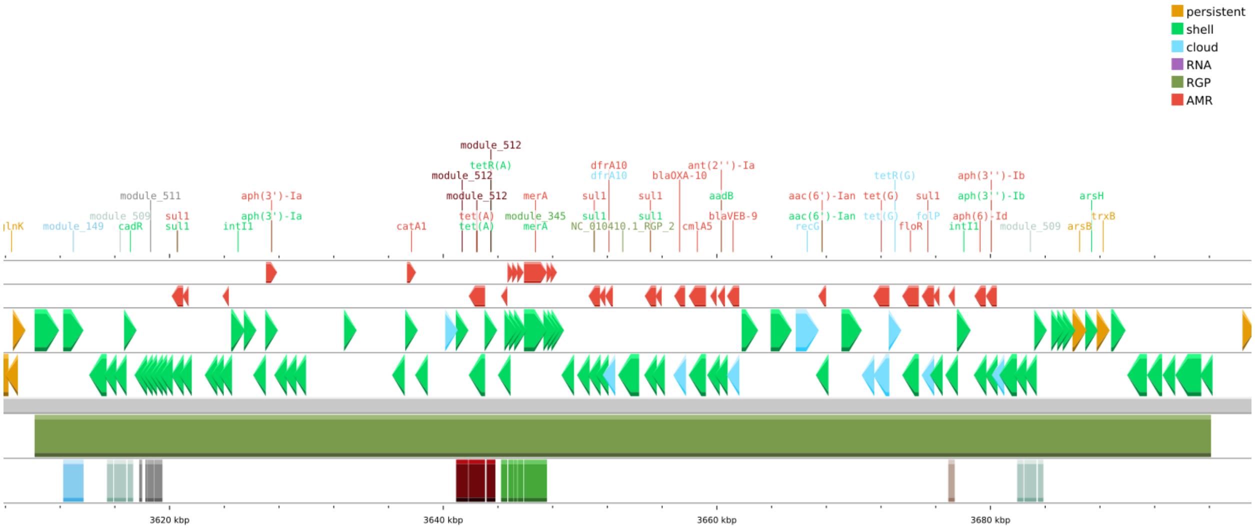


**Figure S6: Genomic organization of the AbaR1 resistance island in *Acinetobacter baumannii* strain AYE (GCF_000069245.1).** CGView genomic map centered on the AbaR1 resistance island, structured from top to bottom as follows: (1) AMR genes: red arrows indicate AMR genes and their transcriptional orientation (2) Genes: coding sequences are represented as arrows oriented according to their transcriptional direction and colored by pangenome partition: *persistent* (orange), *shell* (green) and *cloud* (cyan); non-coding RNAs are represented by purple arrows (3) RGP: The solid olive green rectangle defines the boundaries of the RGP (4) Modules: coding sequences belonging to predicted modules are highlighted by colored boxes. Genomic coordinates (in kbp) are indicated along the x-axis.

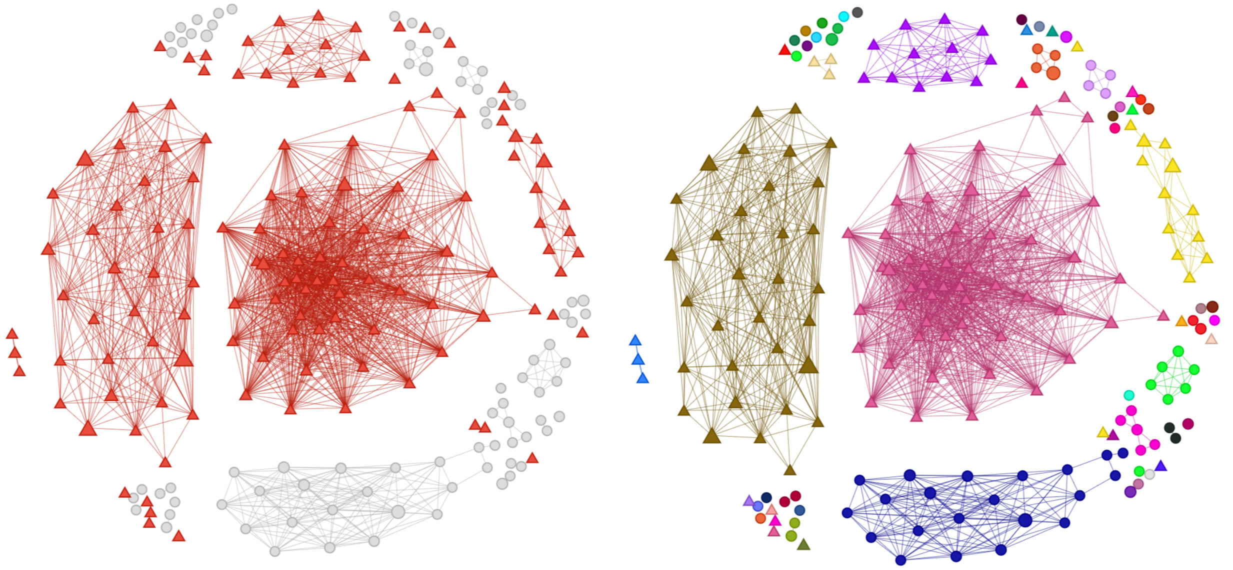

**Figure S7: Network representation of clustered RGPs within the AbaR resistance island in *Acinetobacter baumannii*.** Nodes represent groups of identical RGPs inserted at spot 47, with node size proportional to the number of RGPs. Edges connect RGP groups sharing $\geq 80\%$ gene family content (*max_grr* parameter of the RGP clustering functionality of PPanGGOLiN). Left graph: global network where nodes carrying at least one AMR gene family are depicted as red triangles, and AMR-free RGPs are shown as light-gray circles. Right graph: network partitioning into 56 distinct RGP clusters, with nodes colored according to cluster membership.

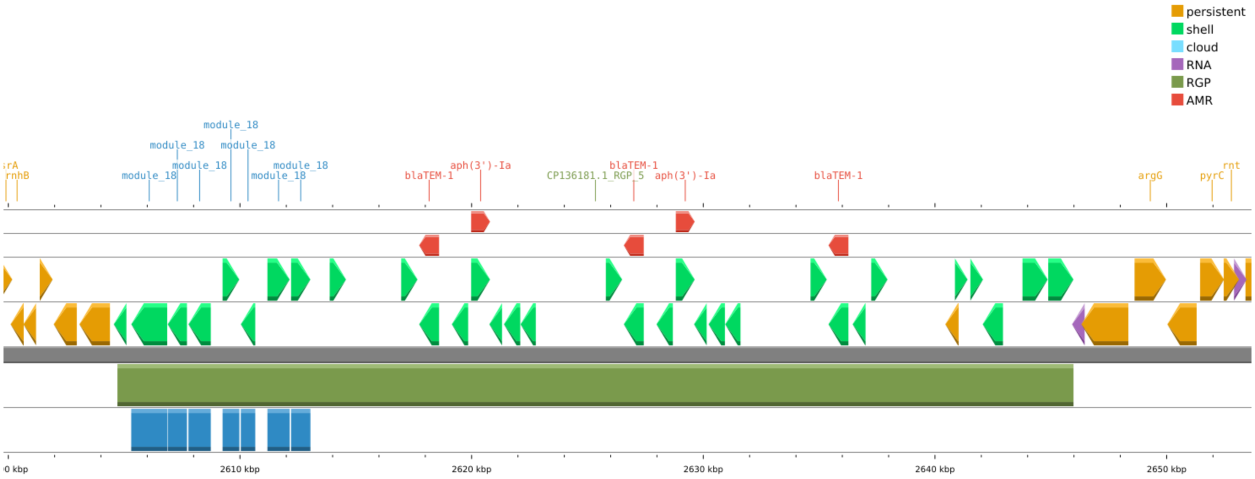


**Figure S8: Genomic organization of the AbGRI2 resistance island in the *Acinetobacter baumannii* canine isolate ABO21-A003 (GCF_033191195.1).** CGView genomic map centered on the AbGRI2 resistance island, structured as described in Figure S6. The RGP carries three copies of *blaTEM-1* (beta-lactamase) and two copies of *aph(3')-Ia* (aminoglycoside phosphotransferase).
